# Contraction-expansion dynamics of reef species in response to sea-level changes

**DOI:** 10.1101/2021.01.06.425656

**Authors:** TB Hoareau, PC Pretorius

**Affiliations:** Department of Biochemistry, Genetics and Microbiology, University of Pretoria, X20, Hatfield 0028, Pretoria, South Africa; Reneco International Wildlife Consultants LLC, Sky Tower, office 3902 and 3903 - Al Reem Island P.O. Box 61741 - Abu Dhabi, United Arab Emirates

**Keywords:** Historical demography, Marine transgression, Paleoecology, Pleistocene, Scleractinia

## Abstract

The contraction-expansion model (CEM) describes the dynamics of species that survived in refugia during the last glacial maximum (LGM) and expanded their range when environmental conditions slowly improved from the Late Glacial through to the Holocene. The CEM has been proposed to describe the dynamics of reef species in response to sea-level fluctuations from a range of disciplines, but genetic inferences rather suggest stable population sizes since the last glacial period. Here, we address this paradox by providing a new model of modern reef development, by assessing the effect of LGM bottlenecks using genetic simulations, and by using a survey of the literature on reef species to compile both estimates of times to expansion and applied rates of molecular evolution. Using previously published radiocarbon dates of core data, we propose a synthetic model for the dynamics of modern coral reefs in the Indo-Pacific region. This model describes both an initiation at 9.9 ka and subsequent development that confirms a strong influence of sea-level fluctuations on reef dynamics. Simulations based on mtDNA datasets showed that pre-LGM genetic signatures of expansion are lost. Recent literature shows that, although genetic expansions of tropical marine species are frequent (>95%), the onset of these expansions is old (median ~110 ka), which indicates that most populations have remained stable since before the LGM. These pre-LGM expansions are explained by the low mutation rates (1.66% changes/site/Myr) known to be inadequate to calibrate time at population level. Specific calibrations should help solve the paradox and generalise the CEM for reef species.

## Introduction

The contraction-expansion model (CEM) has been proposed to describe the demographic response of coral reef species (abundance, distribution, and gene flow) to sea-level fluctuations (Paulay 1990 and 1996; Crandall et al. 2008; Camoin and Webster 2015; Ludt and Rocha 2015). According to this model, species experience a bottleneck when sea levels are at a low stand during glacial periods, while they recover and thrive towards the end of marine transgression during interglacial periods (Hewitt, 2000; Provan and Bennett 2008; Hoareau et al. 2012; Grant 2015). For instance, signatures of erosion are identified during the Last Glacial Maximum (LGM; ~20 ka) when reefs were exposed to weather conditions at low sea level, while significant depositions appeared during the Holocene period when the sea level was rising (Paulay and McEdward 1990). The drop in sea level resulted in up to 95% loss of habitat with only fore reefs left during the LGM (Paulay 1990; Voris 2000), which was then recovered during the Holocene (Paulay 1990; Crandall et al. 2012; Ludt and Rocha 2015). Fossils and genetic evidence also support cases of extinctions occurring during the LGM, while entire habitats were recolonised by reef species during the Holocene (Pizarolli and Montaggioni 1988; Walbran et al. 1989; Paulay 1990; Ludt and Rocha 2015). These LGM conditions most likely resulted in demographic bottlenecks for reef species that survived, while evidence of expansion during the Holocene for a majority of species is clear from the literature (Crandall et al. 2012; Grant 2015; Ludt and Rocha 2015). A consensus therefore seems to be reached on the effect the sea-level fluctuations had on coral reefs, but there are still uncertainties on the timing of reef expansions as it heavily depends on basement substrates (Camoin and Webster 2015).

As historical events can leave a clear signature in the genomes of modern individuals, estimating of genetic history from nucleotide sequences can provide precious insights about the impact of climate changes and anthropogenic factors (Ho and Shapiro 2011). Sequence mismatch analysis (Rogers and Harpending 1992) and skyline plots (Pybus et al. 2000; Drummond et al. 2005) are frameworks that can help give a timeline on past demographic expansions. Applied to reef species, these methods often detect expansions, yet these are so old that population sizes remained constant during the LGM (Hoareau et al. 2013; Delrieu-Trottin et al. 2017). If the timeline of these events is correct, these results suggest that the LGM bottlenecks identified by other disciplines had left no signatures on the genetic polymorphism of these species.

An accurate inference of past demographic histories can only be obtained by applying a molecular clock based on appropriate mutation rates. Testing the validity of the molecular clock is therefore a key step before assessing the robustness of estimates of time to expansion or evaluating the impact of specific environmental factors on the demography. Over the last decade, several studies have shown that molecular clocks based on phylogenetic calibrations provide rates that are too low to produce accurate intraspecific time and demographic parameters (The so-called time-dependency of the rates; Ho et al. 2005; Ho et al. 2011; Grant 2015). Yet, these rates are still dominant in the calibrations of population events. The inflated times based on these rates create a mismatch between historical changes in the environment and the genetic inferences (Ho et al. 2008; Grant 2015; Hoareau 2016). Recent studies have provided ways to account for these biases, but although some have been applied to coral reef species (Crandall et al. 2012; Hoareau 2016), it is difficult to generalise their applications to other tropical species associated with tropical marine coastal ecosystems. New calibrations that can be generalised to reef systems must rely on a strong environmental proxy that reflect the dynamics of species associated to the ecosystem.

This paper explores the validity of the CEM for reef species in the light of the paradox created by the results obtained from genetic inferences. The objectives include 1) creating a synthetic model for the development of modern coral reef (Reef Growth phase RGIII; Montaggioni 2005) in the Indo-Pacific region; 2) evaluating the effect of a LGM bottleneck on past genetic signatures applying inferences of population history on simulated sequence data; 3) compiling times to expansion of reef species obtained from the literature in relation to the development of modern coral reefs and late Pleistocene sea level changes; and 4) compiling rates of molecular evolution applied to these reef species in relation to the challenge associated to the time-dependency of the rates.

## Materials and methods

### Initiation and development of modern coral reefs

We recovered maximum radiometric age of modern and post-LGM submerged coral reefs from the Indo-Pacific region found in Montaggioni (2005) and references herein (Table S1). Coral reef was considered modern if it was part of the reef growth phase RGIII as defined in Montaggioni (2005). Following this same study, we considered the maximum age (age of the reef base) as an approximation of the initiation of each coral reef complex. We excluded Vanuatu’s reef as there is evidence of accretion linked to uplifting (Cabioch et al. 1998) which may bias the analysis.

We identified the transition time between modern and submerged coral reefs using the density function of a logistic regression analysis, with the age of coral reefs as dependent variable and the modern/submerged characteristics as independent variables. We then produced a cumulative curve of the initiation ages of modern coral reef, which provided an approximation of their development during the Holocene period. Montaggioni (2005) used a classification of modern coral reefs into three types: 1) exposed reef crests and flats; 2) semi-exposed to sheltered reef crests and flats; 3) backreef zones.

We used this partitioning to produce habitat-based cumulative curves and explore variability of habitats dynamics. Variation in sea level was represented using data derived from a standard chronology for oxygen isotope records produced by several SPECMAP reference series (Waelbroeck et al. 2002) that are unrelated with coral reef dynamics.

### Effect of LGM bottleneck intensity on earlier genetic signatures

To test the underlying idea that strong bottlenecks can erase genetic diversity, thereby preventing the correct inference of past events, we used FASTSIMCOAL 2 (Excoffier et al. 2013) to simulate different sets of non-recombining sequences of 1000 nucleotides. The simulations were done using demographic scenarios mimicking variations in sea level observed over the last 200 kyrs, but each with varying intensity of bottleneck associated with the LGM. The decrease in *Ne* during the bottleneck is illustrated by the ratio log(*Nebottleneck*/*Neoriginal*) that ranged from 0.2 to 80% of the pre-bottleneck population size, itself set to 500,000 individuals. The mutation rate applied was 2×10^−7^ changes per site per year using a HKY substitution model, which is close to rates obtained from aDNA studies (Ho et al. 2008) or to those based on demographic calibrations (Crandall et al. 2012; Hoareau 2016).

We obtained variations in *Neτ* (with *τ* the generation time) over the time parameter *t* from the simulated sequences applying the extended Bayesian Skyline plot model (EBSP; Heled and Drummond 2008) available in the program BEAST version 2.6.2 (Bouckaert et al. 2019). We applied the HKY model of nucleotide substitution and the mutation rate as used for the simulations. A total of 10,000 genealogies and model parameters were obtained after discarding 10% of samples as burn-in from a total of 110 million iterations (MCMC parameters). We used TRACER v1.7.1 (Rambaut, Drummond, Xie, Baele and Suchard, 2018) to check the quality of the runs by verifying that the effective sample sizes (ESSs) of all parameters were above 200, as suggested by the BEAST guidelines. We recovered the extended Bayesian Skyline Plot and the TMRCA posterior distribution to represent both the population history and the oldest time recovered for the ancestor of the population for illustration of the effect of the intensity of bottleneck.

### Literature survey

To provide an overview of initiation of expansions in reef species and associated mutation rates that were applied, we surveyed the literature published between 2015 and 2020 (up until 22^nd^ September 2020) using the google scholar search engine. We used a procedure described by Grant (2015) who used the following search terms: Bayesian skyline plot, mismatch analysis, mismatch distribution, historical demography, mitochondrial DNA, molecular clock calibration, mutation rate. We included additional search terms (reef*, coral*) to specifically target reef species. We subsequently checked each paper more closely, downloaded the relevant ones, and recovered both the rates of molecular evolution used and the dates of expansion obtained.

## Results

### Synthetic model for the development of modern coral reefs

The transition time between submerged and modern reefs was modelled by the density function of a logistic regression, which provided an estimate for the establishment of modern coral reefs at 9.876 (6.674-13.079) ka (Fig. 1A). The accumulation curve describing the succession of dated coral reefs in the Indo-Pacific shows a sharp increase, especially between 8 and 6 ka. After 6 ka, the accumulation becomes moderate but continues until more recently (Fig. 1B). We observed differences between the types of reef habitats (Fig. 1C), with the most significant pattern being the strong increase of reefs found in backreef zones around 6 ka. Considering the changes in relative sea level derived from oxygen isotope records (Waelbroeck et al. 2002), the Pleistocene-Holocene transition (11.7 ka; Walker et al. 2009) occurred at an average sea-level stand of 55.4 (± 6.5) m below present, the establishment of modern coral reefs (9.876 ka) at 29.8 (16.6-42.6) m, and the transition in reef development observed around 6 ka at 4.2 (± 1.0) m.

**Fig. 1.**
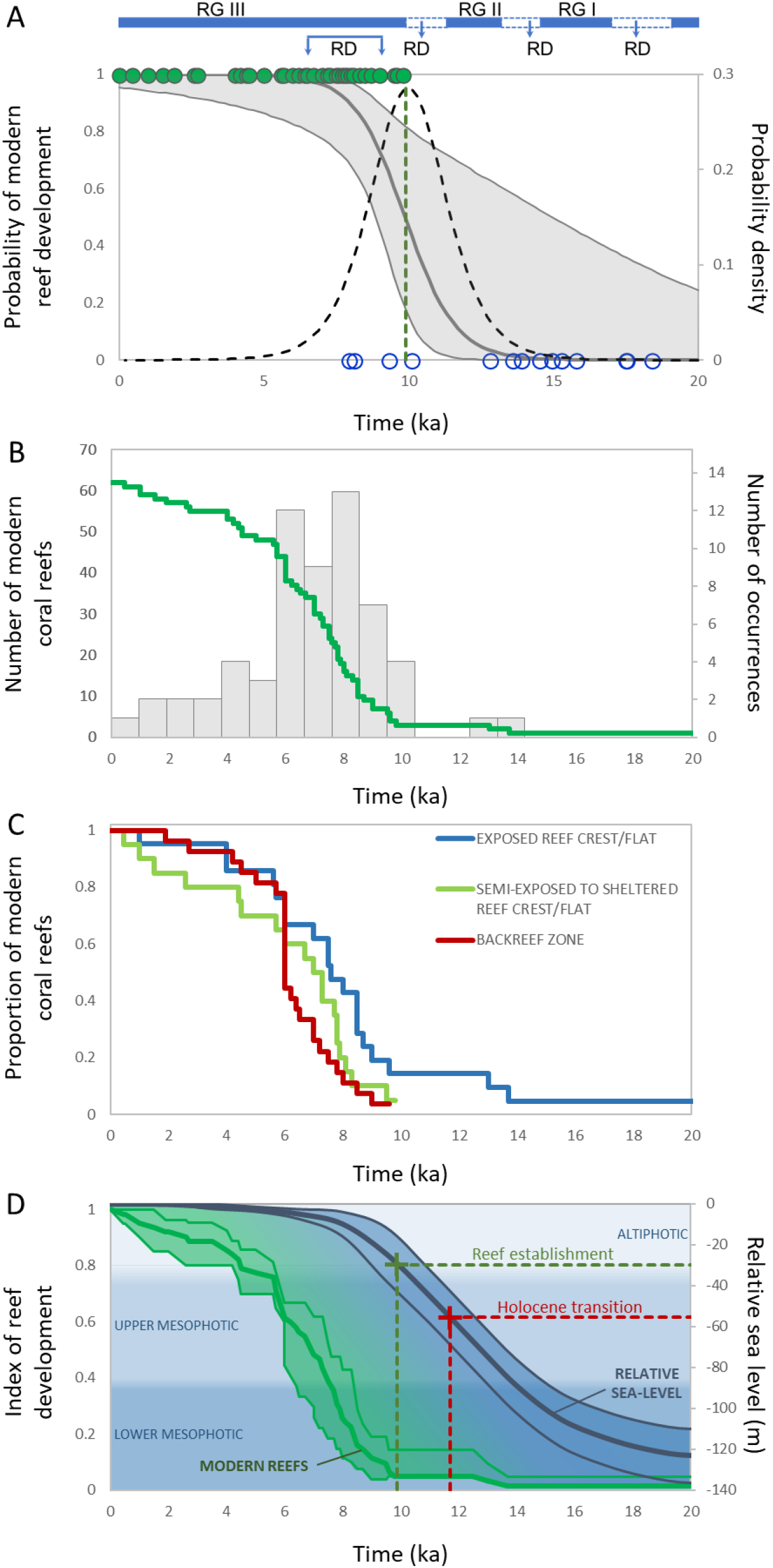
Responses of coral reefs to quaternary sea-level changes in the Indo-Pacific region. **A)** Onset of the development of modern coral reefs during the Holocene illustrated by the density function (dotted black curve) of a logistic regression (Wald test: χ^2^ = 5.8, *P* = 0.016) between post-LGM submerged reefs (open dots) and corals reefs that survived (full dots), based on previously published radiometric ages (Montaggioni 2005, and references herein). RGI, RGII, RGIII are Reef Growth periods and RD are events that are considered non-constructional or reef drowning (Montaggioni 2005). The transition time between submerged and modern reefs occurred at 9.876 (6.674-13.079) ka (green dotted vertical line), which coincides with the start of RGIII at 10 ka. The cumulative distribution function of the logistic regression is also shown in grey for illustration purpose. **B)** Frequency distribution and cumulative curve of the ages of all modern coral reefs. **C)** Cumulative curves of different types of reef habitats assumed to have different dynamics over time (Montaggioni 2005): exposed reef crests and flats developed first, then semi-exposed to sheltered reef crests and flats, followed by backreef zones that have a strong increase around 6 ka towards the end of the rise in relative sea level (4.2 ± 1.0 m). **D)** Synthetic model illustrating the establishment and accumulation of modern coral reefs in the Indo-Pacific region, the changes in relative sea level (Waelbroek et al. 2002), the Holocene transition at 11.7 ka (Walters et al. 2009), the establishment of modern reef, as well as the reef faunal zonation (Baldwin et al. 2018). The establishment of modern reefs corresponds to an average sea-level stand of 29.8 ± 6.5 m below current sea-level.

### Effect of LGM bottleneck intensity on earlier genetic signatures

Assuming a demography strictly influenced by the relative sea level, and considering different intensities in population changes, the simulation study indicates that the most recent expansions associated with the post-LGM deglaciation are the only ones that can be detected (Fig. 2A and 2B). The genetic inferences provide an appropriate representation of the relative intensity of simulated expansions during the deglaciation. Moreover, the distribution of posterior probabilities of TMRCAs for these inferences (proxy for the depth of genetic history) decrease with higher intensities of bottlenecks (Fig. 2C), and this is supported by the inverse correlation between the two parameters (Fig. 2D).

**Fig. 2.**
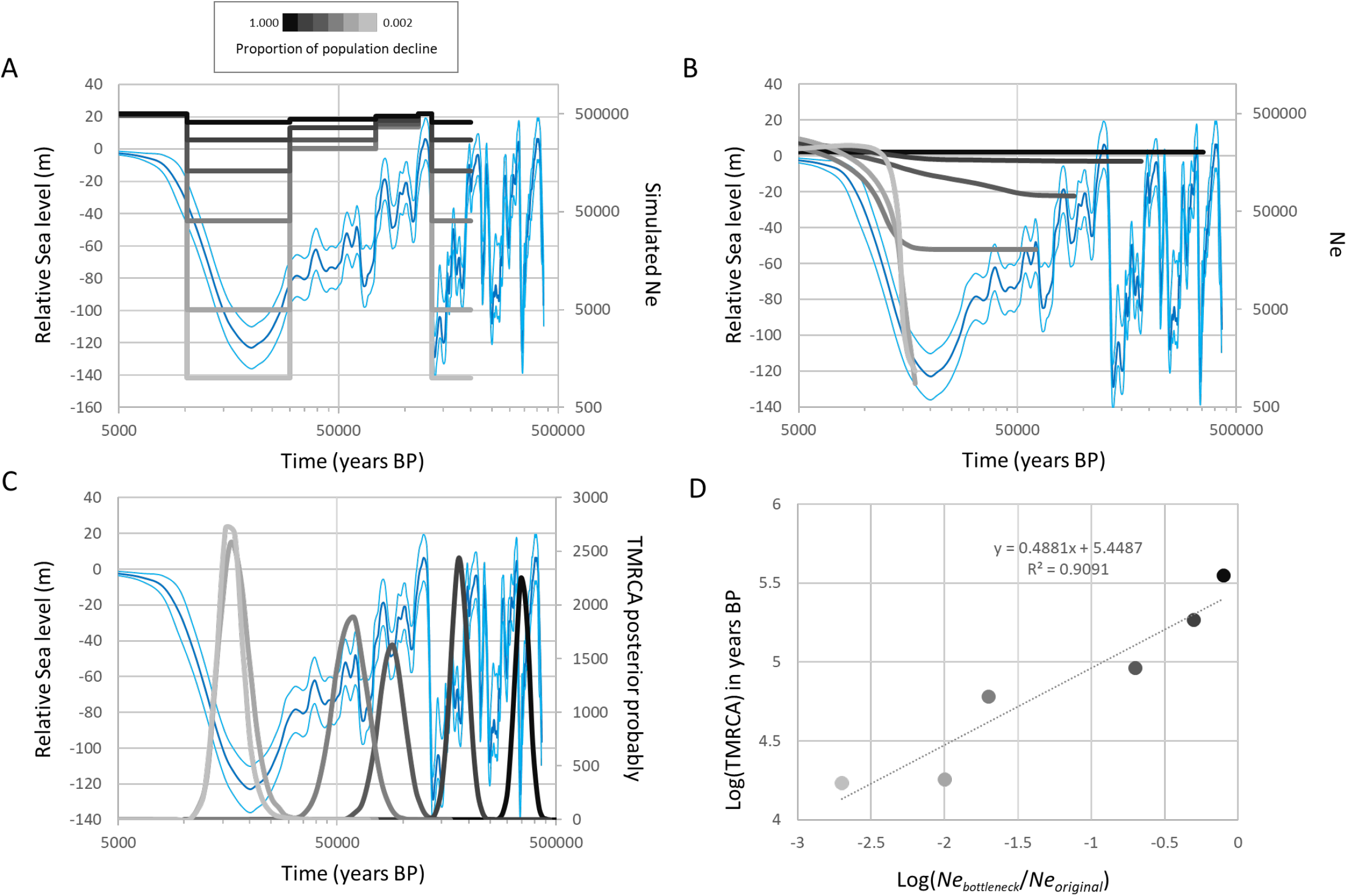
Extended Bayesian Skyline Plot model (EBSP model) applied to simulated nucleotide sequences to illustrate the effect of different levels of population contractions during the LGM on demographic reconstructions. **A)** Demographic scenarios used to simulate the haploid nucleotide sequences overlaid to the relative sea level over the last 500 ka (Waelbroek et al. 2002). **B)** Calibrated reconstruction for the different scenarios. **C)** Posterior probabilities of calibrated TMRCA obtained for the different scenarios. **D)** Scatter plot with linear fit showing the logarithms of TMRCA as a function of the intensity of population contraction Log (*Ne_bottleneck_*/*Ne_original_*). The regression model is *y* = 5.4487 + 0.4881*x*, Pearson *R* = 0.9535, *P* < 0.0032.

### Timeframes of demographic expansion for reef species and associated rates of molecular evolution

The literature survey from 2015-2020 resulted in a selection of 101 papers that provided 431 instances with demographic status that included 413 cases of expansions (95.8%), while the rest was considered to have constant sizes. Among these, a total of 401 provided a date for the expansion, with 358 (89%) pre-dating 20 ka (LGM) and 43 (11%) post-dating the LGM. A total of 102 estimates was obtained for the rates of molecular evolution. The terminology for the rate used varied and included substitution rates, mutation rates, lineage rates and divergence rates among others. Most of the rates were low with a median value at 1.66×10^−8^ (95% percentile: 0.14×10^−8^-5.00×10^−7^) changes per site per year.

## Discussion

### Contraction-expansion dynamics of coral reefs in response to sea-level fluctuations

The new synthetic model of reef development (Fig. 1) is based on radiocarbon dating from various reefs throughout the Indo-Pacific region and provides precious information about the dynamics of modern coral reefs. This model shows that the initiation happened around 9.9 ka, which coincides with the beginning of the reef generation RGIII that marks the accretion of modern coral reefs between 10–7 ka (Montaggioni 1988, 2005; Camoin and Webster 2015). After this transition, the number of drowned reefs significantly decreases (Fig. 1A), indicating that coral reefs were, for the first time since the LGM, in keep-up mode of growth with marine transgression (Camoin and Webster 2015). Around 10 ka, sea levels ranged between 20 and 40 m below present level (Fig. 1D; Waelbroek et al. 2002), which encompasses the 30–40 m below present depth range that is thought to coincides with the colonization of inner shelf substrates and reefs (Carter and Johnson 1986). This is confirmed by the thickness of modern reefs that reach up to 30 m (Montaggioni 2005). These sea level ranges also coincide with the lower limits of the altiphotic zone (40 m) that has recently been proposed to describe the shallowest coral reef faunal zone characterized by high light levels (Fig. 1C; Baldwin et al. 2018). This is in support of previous findings that showed that the initiation of modern coral reefs was controlled by the pace of increase in sea level (Montaggioni 2005; Camoin and Webster 2015) and other environmental factors including light conditions (Laverick et al. 2019).

The model further shows that the number of coral reefs increased throughout the Indo-Pacific region after the initiation period until around 6 ka, when the overall increase in sea level decelerated to become null towards the present (Fig. 1C). Most types of habitats have the same trends as the general pattern of reef development, except the backreef zone that has a sharp increase around 6 ka. This mid-Holocene increase in backreef habitats has previously been observed in several parts of the Indo-Pacific region (Camoin and Webster 2015), including on the Great Barrier Reef (Sanborn et al. 2020). Around this time, the sea level reached its present position (Fig. 1D; Waelbroek et al. 2002), which may coincide with specific reef initiation following sudden substrate flooding in backreef zones. Although previous studies have shown that the dynamics of coral reefs is first controlled by basement substrates (Camoin and Webster 2015), the model presented here confirms a strong influence of sea-level fluctuations.

### Evaluating the contraction-expansion model in reef species using genetic inferences

To evaluate the potential role sea levels have had on past demography of coral reef species, we compared the effect of varying intensities of LGM bottlenecks on demographic inferences using a simulation study. The inferences based on the EBSP model represent fair reconstructions of the simulated population histories, which is in line with recent studies (Hoareau 2016, 2020; Pretorius and Hoareau 2020) and confirms the robustness of the analysis. Our results show an absence of expansion older than the LGM (Fig. 2), indicating that LGM bottlenecks erase old signatures of expansion from a typical mtDNA dataset that is traditionally used in phylogeography. This effect is further illustrated by the relationship between intensity of bottlenecks and TMRCAs (Fig. 2C), the latter being often considered a proxy for the depth of history recovered from genetic data. By removing historical genetic polymorphism, declines in population size prevent the reconstruction of pre-bottleneck expansion from mtDNA genealogies, which support previous works (Grant and Cheng 2012; Grant 2015). As a result, no expansion is expected to be recovered from sequence data obtained from most reef species since they have experienced intense bottlenecks during the LGM.

The literature survey indicates that a large majority of species inhabiting reef ecosystems experienced a demographic expansion (93%). However, unlike the expectations under the contraction-expansion model described above, the demographic expansions of most reef species are inferred to be much older than the LGM, with onset of expansions varying widely and going up to millions of years ago (Fig. 3a).

**Fig. 3.**
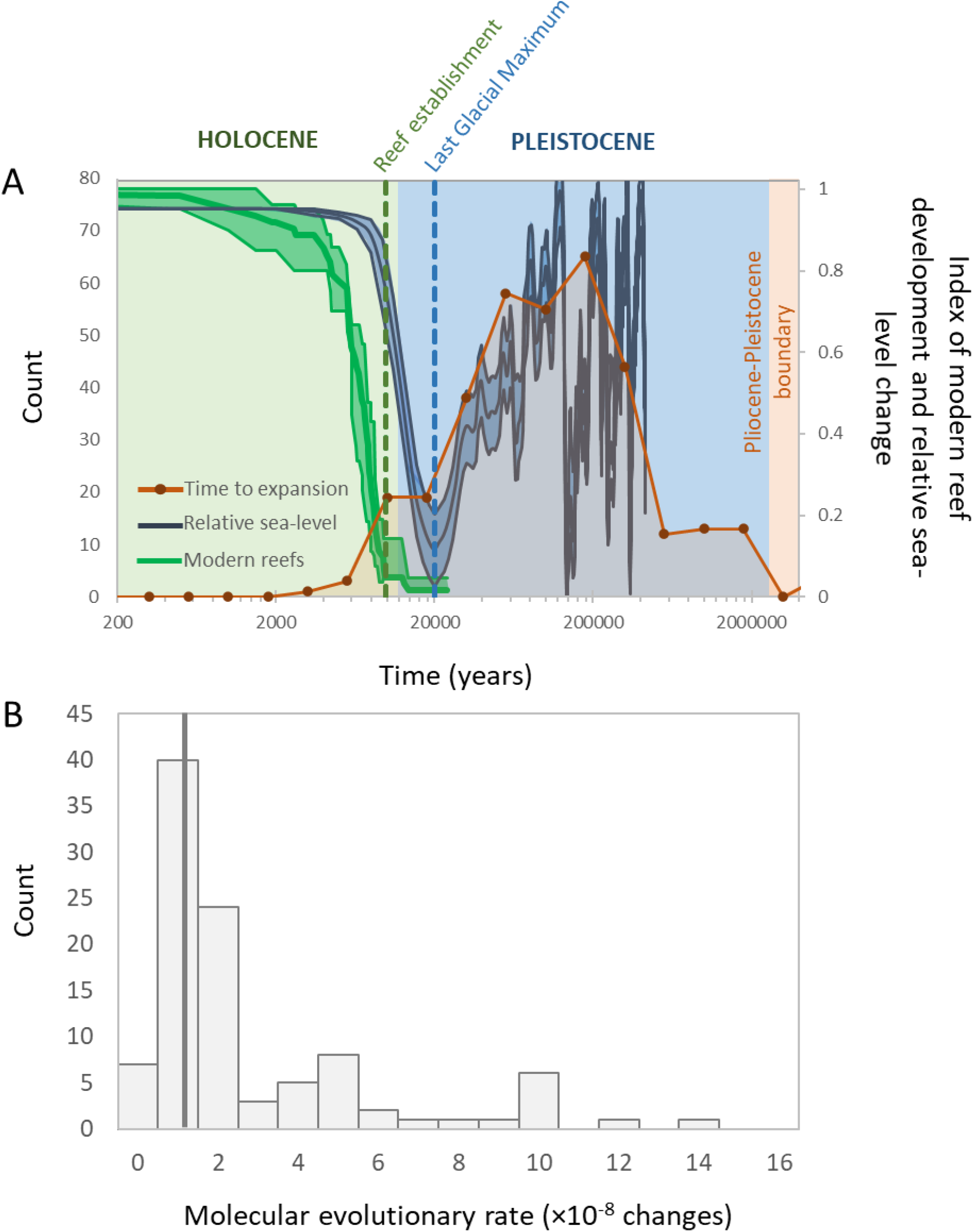
Results of the literature survey based on articles published between 2015 and 2020 and illustrating that inferred expansion times are much older than the LGM for reef species. A) Distribution of estimates of expansion times over the last four million years (*t* = 110 (95% percentile: 11-1712) ka; n = 401) and associated relative sea level and modern reef development. B) Molecular evolutionary rates recovered (n = 102) that were used to calibrate the timing of demographic expansion in reef species. The median value is 1.66×10^−8^ (95% percentile: 0.14×10^−8^-50.00×10^−8^) changes per site per year.

Such old expansions imply that effective population sizes stay constant throughout the LGM, or even over several cycles of glacial and interglacial periods, which is highly unlikely given the strong influence of sea level fluctuations on reef species. Moreover, these dates are very scattered even though the species are found in the same habitats and most share the same life cycle. A recent study has suggested that expansions older than the LGM can be explained by ancient colonization and speciation processes (Delrieu-Trottin et al. 2017). These ideas go against ecological expectations and the evidence of post-LGM expansion gathered from geomorphological studies, fossil data or habitat reconstructions (Paulay 1990; Crandall et al. 2012; Camoin and Webster 2015). Genetic inferences suggesting old and conspicuous increase in population size over several glacial periods therefore create a paradox that raises the question of the validity of the timeline provided by these analyses.

In genetic inferences, timing of events relies on the calibration of the molecular clock using appropriate mutation rates. Over the past decade, concerns have been raised about the accuracy of substitution rates derived from old calibrations (>1 Ma) that have been the main source of calibration to date events at the population level (Ho et al. 2005; Ho et al. 2011). The substitution rates used are often too low, which create inflation of time and population parameters (Ho et al. 2008; Ho et al. 2011; Crandall et al. 2012; Hoareau 2016). As expected, our literature survey show that the same pattern emerges for coral reef species, with applied rates that are very low around 1.66×10^−8^ changes per site per year (Fig. 3). This is an order of magnitude lower than mutation rates obtained from ancient DNA (Ho et al. 2005) and lower than rates obtained from methods based on expansion dating applied to reef species (Crandall et al. 2012; Hoareau 2016).

A previous study specifically evaluated the contraction-expansion model for the temperate marine biota from the east Pacific (Marko et al. 2010), but all calibrations were based on old events which could lead to invalidate the original hypothesis. Other genetic studies, however, have brought evidence in support of the contraction-expansion model in marine species (Marino et al. 2011; Crandall et al. 2012; Hoareau et al. 2012). We argue that reef species follow a contraction-expansion model, and that the paradox observed between genetic and other evidence is linked to inadequate mutation rates used. The new calibration method specifically developed for reef species and based on the model described by Pretorius and Hoareau (2020) will help gain further insights on factors that drive the population history of tropical species.

## Acknowledgements

The authors thank Philippe Borsa, Wei-Jen Chen and Cecile Fauvelot from the DHEEP research project for helpful discussions on the demographic reconstruction of reef species. The authors benefitted from the technical support and the bioinformatic resources of the Centre for Bioinformatics and Computational Biology of the University of Pretoria and the Centre for High Performance Computing (CHPC) from the South African Department of Science and Technology (DST). TB Hoareau benefitted from the University of Pretoria’s research fellowship programme and the Research and Innovation Support of University of Pretoria.

